# Testing the perilesional neuroplastic recruitment hypothesis in aphasia

**DOI:** 10.1101/2021.08.06.455431

**Authors:** Andrew T. DeMarco, Candace M. van der Stelt, Sachi Paul, Elizabeth Dvorak, Elizabeth Lacey, Sarah Snider, Peter E. Turkeltaub

**Affiliations:** Department of Rehabilitation Medicine, Georgetown University, Washington, DC, 20057, United States; Department of Neurology, Georgetown University, Washington, DC, 20057, United States; MedStar National Rehabilitation Hospital, Washington, DC, 20010, United States

## Abstract

**Objective:** A prominent theory proposes that neuroplastic recruitment of perilesional tissue supports aphasia recovery, especially when language-capable cortex is spared by smaller lesions. This theory has rarely been tested directly, and findings have been inconclusive. Here, we test the perilesional plasticity hypothesis using two fMRI tasks in two groups of stroke survivors.

**Methods:** Two cohorts totaling 84 chronic left-hemisphere stroke survivors with prior aphasia diagnosis, and 80 control participants underwent fMRI using either a naming task or a reliable semantic decision task. Individualized perilesional tissue was defined by dilating anatomical lesions, and language regions were defined using meta-analyses. Mixed modeling examined differences in activity between groups. Relationships with lesion size and aphasia severity were examined.

**Results:** Stroke survivors exhibited reduced activity in perilesional language tissue relative to controls in both tasks. Although a few cortical regions exhibited greater activity irrespective of distance from the lesion, or only when distant from the lesion, no regions exhibited increased activity only when near the lesion. Larger lesions were associated with reduced language activity irrespective of distance from the lesion. Using the reliable fMRI task, reduced language activity related to aphasia severity independent of lesion size.

**Interpretation:** We find no evidence for neuroplastic recruitment of perilesional tissue in aphasia beyond its typical role in language. Rather, our findings are consistent with alternative hypotheses that left-hemisphere activation changes during recovery relate to normalization of language network dysfunction and possibly recruitment of alternate cortical processors. These findings clarify left-hemisphere neuroplastic mechanisms supporting language recovery after stroke.

**Summary for Social Media If Accepted:** *Twitter handle:* @crlgeorgetown

*What is the current knowledge on the topic?:* After left-hemisphere stroke, many individuals experience long-term language impairment (aphasia) while others recover their communication abilities. Although there are several hypotheses concerning the kind of brain neuroplasticity that allows some individuals to recover, these mechanisms are not understood in aphasia.

*What question did this study address?:* This study tested the perilesional plasticity hypothesis as it relates to aphasia recovery. This predominant theory posits that tissue around the stroke lesion boundary becomes recruited to support recovered language function in post-stroke aphasia.

*What does this study add to our knowledge?:* This study clarifies the mechanisms of neuroplasticity in stroke aphasia recovery. The results are not consistent with the conventional perilesional plasticity hypothesis, but rather favor an interpretation that recovery is supported by normalization of language network dysfunction and possibly recruitment of alternate brain regions

*How might this potentially impact on the practice of neurology?:* These conclusions will give practicing neurologists a better understanding of how the brain recovers from aphasia after stroke.

## Introduction

Stroke is a leading cause of permanent disability, and sequelae are partially determined by lesion size and location. However, an important driver of recovery is thought to be neural reorganization in residual tissue beyond the lesion boundaries (Grefkes & Fink, 2020). A mechanistic account of this plasticity is necessary to make progress in aphasia neurorehabilitation, and several mechanisms have been proposed (Stefaniak et al., 2020; Turkeltaub, 2019).

Among the proposed mechanisms is the perilesional plasticity hypothesis, which emphasizes tissue immediately surrounding the lesion, where animal studies have both observed dysfunction (Neumann-Haefelin & Witte, 2000; van der Zijden et al., 2008) and suggested collateral axonal sprouting and synaptogenesis may support functional recovery (Nudo, 1999; Stroemer et al., 1995). Motor stroke recovery, in particular, appears to rely on functional take-over by perilesional sensorimotor (Teasell et al., 2005) or primary motor cortices (Jaillard et al., 2005; Xerri et al., 1998).

These findings have informed models of aphasia recovery, which stipulate that when language tissue is damaged, alternative perilesional processors may become recruited to support outcomes, especially around small lesions (Heiss & Thiel, 2006; Thompson & den Ouden, 2008). Treatment studies provide the strongest evidence for perilesional recruitment, finding increased activity after treatment that relates to gains in performance (Fridriksson et al., 2012; Meinzer et al., 2008). However, because these studies have not compared the activation directly to control subjects, they cannot clearly establish whether treatment-related increases in perilesional activity represent either neuroplastic recruitment of new tissue for language or supranormal recruitment of typical language regions due to plasticity.

Alternatively, treatment-related increases in perilesional activity may reflect normalization of function in language tissue that is dysfunctional due to network effects of the nearby lesion. Studies of spontaneous stroke aphasia recovery have found an acute reduction in left-hemisphere language activity, followed by subacute supranormal activity and a chronic normalization of activity associated with good outcomes (Saur, 2006). While increased perilesional activity was associated with better performance, the activity did not exceed that of controls (Stockert et al., 2020). These findings suggest that lesions cause language network dysfunction, and that recovery is supported by normalization of activity rather than by recruitment of new perilesional tissue into the language network.

Here, we test predictions of the perilesional plasticity hypothesis, contrasting activity elicited by two independent language tasks in a large cohort of left-hemisphere stroke survivors and matched controls. We predicted that neuroplastic recruitment would result in supranormal perilesional task-related activity. We tested two related predictions: that recruitment (i) might only occur within, or proximal to, language tissue and (ii) might only be evident around small lesions. We also examine whether the effect only occurs in specific brain regions. Finally, we consider alternative hypotheses explaining left-hemisphere activity observed in prior aphasia studies, namely that this activity is either (1) residual activity in the language network not resulting from recruitment of new tissue, or (2) that it represents recruitment of alternate processors irrespective of proximity to the lesion.

## Methods

### Participants

Participants included 84 survivors of left-hemisphere stroke with a history of aphasia, and 80 controls (**Table 1**). Participants were recruited in the Washington, DC area for a clinical tDCS study (naming task data, Doris Duke Charitable Foundation Grant 2012062, 2013-2018), and an ongoing cross-sectional study of aphasia outcomes (semantic decision task, NIDCD R01DC014960, 2018-2020). Between the studies, the MRI scanner was upgraded (see details below). Aside from stroke, participants with aphasia had no other history of significant psychiatric or neurological condition. All participants provided informed consent in accordance with the Georgetown University Institutional Review Board.

**Table 1.** Participant demographics.

| Task | Group | Sex (F / M) | Age Mean (SD) [range] | Stroke Chronicity (Months) Mean (SD) [range] | WAB AQ Mean (SD) [range] |
| --- | --- | --- | --- | --- | --- |
| Naming | Controls | 19 / 21 | 58.5 (12.5) [26-80] | N/A | N/A |
|  | Aphasia | 19 / 32 | 60.1 (9.8) [38-78] | 65.7 (58.8) [1.6 - 283.1] | 71.2 (24.6) [13.8 - 99.8] |
| Semantic Decision | Controls | 20 / 19 | 13.0 (13) [31-82] | N/A | N/A |
|  | Aphasia | 13 / 18 | 61.3 (10.3) [43 - 81] | 40.6 (53.9) [2.0 - 211.4] | 82.2 (18.1) [32.2 - 99] |

### Image Acquisition

Sessions for the naming task were conducted on a Siemens 3T Trio. Sequences included a high-resolution T1-weighted scan (TR = 1900, TE = 2.52, 176 0.9mm sagittal slices, FOV = 240, matrix = 256×256, FA = 9°), a T2-weighted scan (TR = 3200, TE = 45, 176 1.25mm sagittal slices, FOV = 240, matrix = 192×192, FA = 120°), and the BOLD T2*-weighted scan (TR = 2500, TE = 30, 47 3.2mm axial slices, FOV = 204, matrix = 64×64, FA = 90°) consisting of 168 volumes and lasting 6:00.

Sessions for the semantic decision task were conducted on a Siemens 3T Trio. Sequences included a high-resolution T1-weighted scan (TR = 1900, TE = 2.98, 176 1mm sagittal slices, FOV = 256, matrix = 256×256, FA = 9°, SMS = 4), a T2-weighted FLAIR scan (TR = 5000, TE = 38.2, 192 1mm sagittal slices, FOV = 256, matrix = 256×256, FA = 120°), and a BOLD T2*-weighted scan (TR = 794ms, 48 2.6mm slices with 10% gap, 2.9mm voxels, FOV = 211mm, matrix = 74×74, FA = 50°, SMS = 4) consisting of 504 volumes lasting 6:40.

### Lesion segmentation and coregistration

Lesion masks were manually segmented on each participant’s MPRAGE and T2/FLAIR images using ITK-SNAP software (Yushkevich et al., 2006; http://www.itksnap.org/) by author P.E.T **(Fig 1)**.

**Figure 1.**
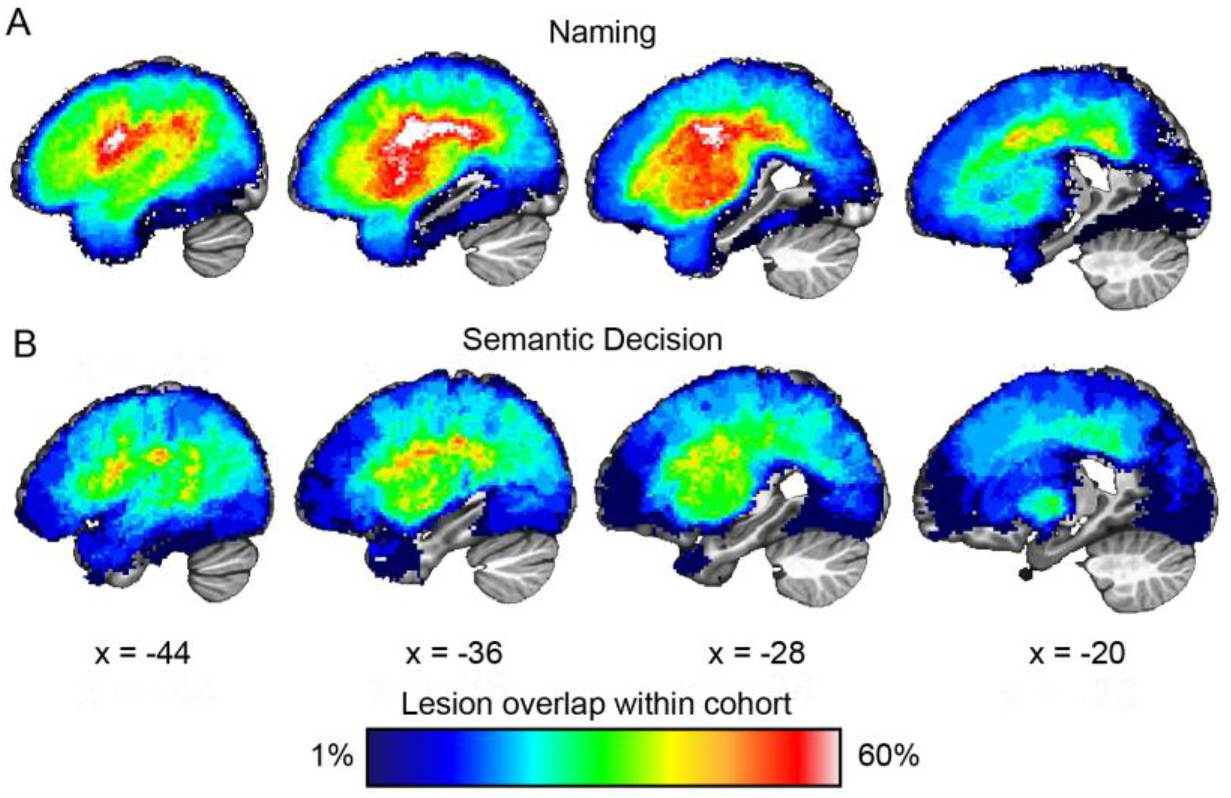
Serial sagittal slices through the left hemisphere of both cohorts showing the overlap of anatomical lesion tracings. Lesion overlap for the cohort who underwent (A, top) the naming task and (B, bottom) the semantic decision task. Percent of lesion overlap within each cohort is indicated by the spectrum color.

### Perilesional tissue definition

We utilized a dilation model of perilesional tissue (Fridriksson et al., 2012) in which perilesional tissue is defined as a shell falling outside of each individual’s anatomical lesion tracing, implemented in MATLAB 2020b using the *imdilate* function (MathWorks, Natick, MA). For brainwide questions, we evaluated 4mm-thick shells spanning 0-4mm, 4-8mm, 8-12mm, and 12-16mm from the lesion boundary. We examined a range of distances from the lesion boundary based on prior work demonstrating reduced perfusion, which may affect task-related BOLD fMRI signal, up to 8mm from the lesion boundary (Richardson et al., 2011). Our further analyses considered perilesional tissue as a single slab between 4-16mm from the lesion boundary. For these slab-based analyses, we discounted voxels immediately neighboring the lesion due to possible partial volume effects (e.g. Stockert et al., 2020). All analyses were restricted to left-hemisphere tissue falling within a standard SPM12 gray-matter tissue probability mask thresholded at >10% likelihood.

### Functional language mapping procedures

Each participant underwent functional language mapping using one of two fMRI tasks. The first task was a common spoken picture-naming task, described in detail in (Skipper- Kallal et al., 2017). The second task was a reliable, adaptive semantic decision task validated in people with aphasia, described in detail in (Wilson et al., 2018). All participants performed greater than chance in the semantic decision task language condition (one-sided binomial test, *P* < .05).

### Image preprocessing and statistical analysis

For both tasks, standard preprocessing was performed in AFNI (Cox, 1996), including slice timing correction, realignment for head motion, despiking, smoothing with a 5mm FWHM kernel, temporal high-pass filtering at 0.01 Hz, and detrending. A whole-brain GLM was estimated using the *fmrilm* function from FMRISTAT (Worsley et al., 2002), with covariates including the time-course of a white matter and CSF seed, and the six head-motion parameters not convolved with the HRF. Each of the 32 naming task trials was modeled using three event types, convolved with the HRF (covert speech period (7.5-9s), overt speech period (5.5s), and fixation (15s)). The contrast of interest was an average of covert and overt greater than fixation [.5 .5 −1]. The semantic decision task was modeled using two alternating boxcar functions (corresponding to the language and control conditions), convolved with the HRF. The contrast of interest was semantic greater than control [1 −1]. Resulting SPMs were then warped to MNI space based on the transformation estimated from the MPRAGE. Finally, images were resliced to 2 mm isotropic voxels.

### Independent task-specific functional definition of brain tissue

To avoid circularity in selecting regions that we planned to analyze (Kriegeskorte et al., 2009), we independently defined language cortex based on meta-analytic results from task-relevant Neurosynth queries (Yarkoni et al., 2011). For the naming task, we operationalized language-cortex (**Fig 2**, C, F, red) as voxels falling within a mask of the result of the search term “speech production” (association test *Z* > 6.0, FDR = .01). We applied the same procedure for the semantic decision task using the search term “language.”

**Figure 2.**
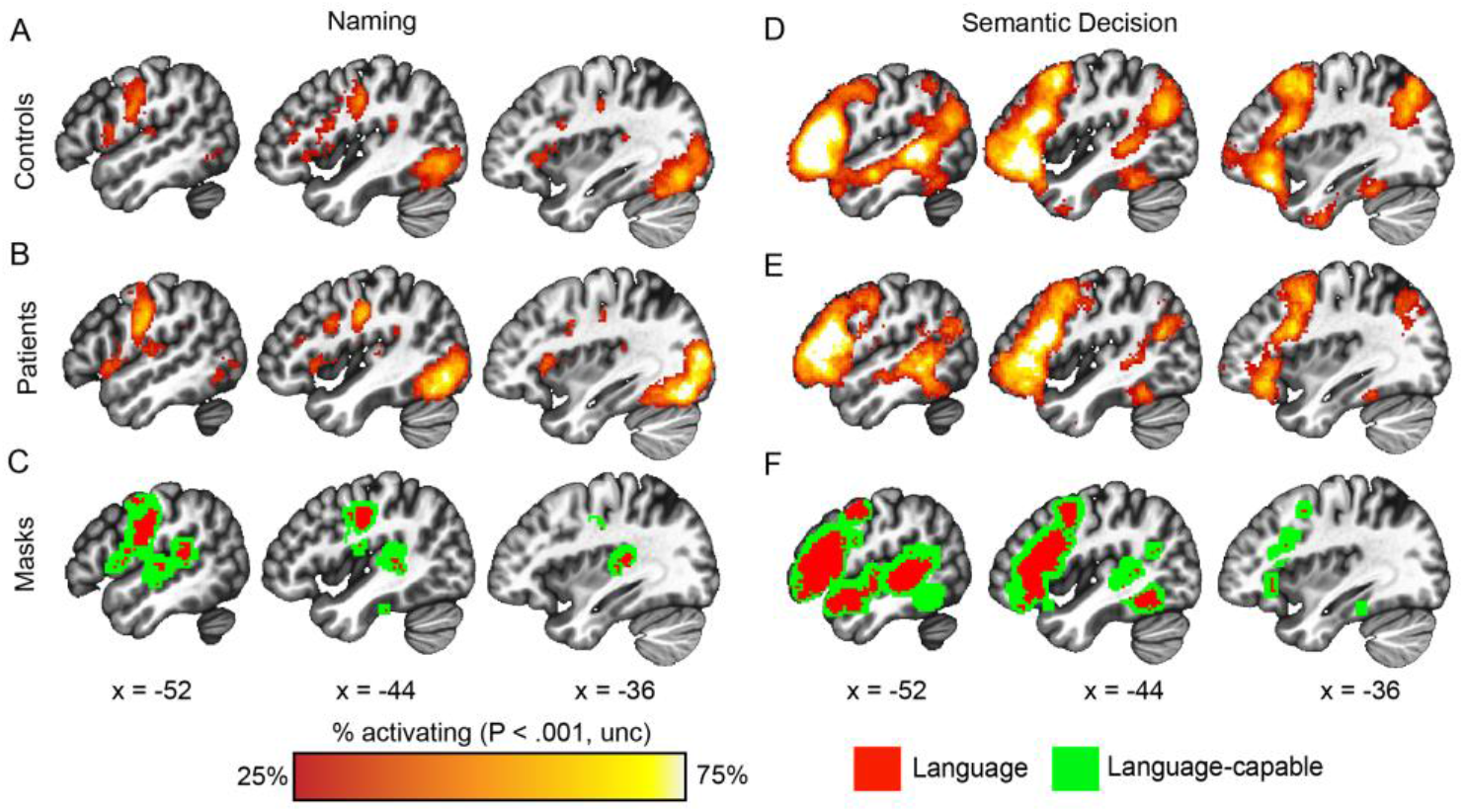
Serial sagittal slices show regions consistently activated across (A, D) the control cohorts and (B, E) aphasia cohorts for the naming task (left) and semantic decision task (right). Percent of each cohort activating is shown as a conjunction of the individuals in each cohort thresholded voxelwise *P* < .001, uncorrected. The bottom row (C, F) shows the extent of independently-defined meta-analytic masks defining task-specific language cortex (red) and language-capable cortex (green).

We operationalized language-capable cortex (**Fig 2**, C, F, green) as an 8mm shell dilated around each task-specific language mask. Finally, we operationalized non-language cortex as voxels falling outside of both language and language-capable masks. Regional analyses were conducted in parcels of a 134 segment atlas (Shen et al., 2013).

### Is perilesional activity different in people with aphasia than in controls?

For each person with aphasia, non-lesioned tissue was first characterized brainwide for each functional tissue category (language, language-capable, non-language) at each of five distances from the lesion boundary (0-4mm, 4-8mm, 8-12mm, 12-16mm, >16mm). For each individual’s masks, we calculated average activation for both that person and for each control subject, applying the person with aphasia’s mask to each control subject in order to compare equivalent tissue. In this way, we generated a set of individualized control values, specific to each person with aphasia, which excludes any contribution from tissue lesioned in that individual.

Then linear mixed effects modeling was used to compare activity in people with aphasia to the controls’ activity while accounting for lesion differences across individuals. The model was repeated for each functional tissue category at each shell distance. The model was specified with a fixed effect of group (aphasia vs control) and random effects of participant and the lesion mask applied to the data (to account for random effects associated with the lesion masks applied to both groups).

For the regional analysis, we calculated a voxelwise intersection of each mask with the atlas to obtain the relevant voxels falling within each region. We then consider effects separately for when a region is near a lesion boundary (4-16mm) and far from the lesion (>16mm). Regions were only examined if at least five people with aphasia had perilesional tissue within its mask. The regional analyses were corrected for multiple comparisons based on false-discovery rate (FDR) at *P* < .05 (Benjamini & Hochberg, 1995).

### Does perilesional recruitment depend on lesion size?

To measure the relationship between perilesional activity and lesion size, we correlated activity and lesion size within each of the three functional tissue types, both in the vicinity of the lesion (4-16mm from the lesion boundary) and distant from the lesion (>16mm from the lesion boundary). For language tissue, we used the linear mixed effects model to test whether individuals with small lesions (<50cc) or large lesions (>100cc) exhibited abnormal activity, relative to controls, in the vicinity of the lesion (4-16mm) or distant from the lesion (>16mm).

### Does activation predict behavioral impairment, independent of lesion size?

We use a semi-partial Spearman correlation (two-tailed) to test whether activity in language regions related to degree of behavioral impairment, independent of lesion size. We focus on a general measure of aphasia severity, the Aphasia Quotient (AQ) from the Western Aphasia Battery (WAB) (Kertesz, 2007). For the naming task, we also examine the relationship between activity and the Naming & Word Finding Subtest from the WAB.

## Results

### Task activation and convergence with tissue masks

Within the left hemisphere, the naming task reliably activated ventral premotor and motor cortex, as well as superior temporal cortex, which is highly consistent with the meta-analysis results (**Fig 2A-C**). The task also reliably activated inferior occipital cortex, likely relating to the visual presentation of the picture stimuli.

We also observed a high degree of consistency between the activation for the semantic decision task and the meta-analytic mask (**Fig 2D-F**). Specifically, participants most reliably activated left inferior frontal cortex, premotor cortex, both anterior superior temporal gyrus and posterior superior temporal lobe, and fusiform gyrus. These results are also highly consistent with previously reported activation using this task (Wilson et al., 2018).

### Perilesional tissue exhibits reduced activity

We first tested whether people with aphasia exhibit perilesional recruitment at various distances from the lesion boundary and within different functional tissue-types. In the naming task **(Fig 3A-C, Table 2)**, language cortex, language-capable cortex, and non-language cortex all exhibited a significant reduction in task-related activation relative to controls, immediately adjacent (0-4 mm) to the lesion boundary. The reduction was evident in both language cortex and language-capable cortex out to 12 mm from the lesion boundary (*P* < .01). In the semantic decision task **(Fig 3D-F, Table 2)**, language cortex exhibited a selective reduction in activation immediately adjacent (0-4 mm) to the lesion boundary (*P* < .01). Although language-capable cortex showed numerically reduced activation, this did not reach significance. Activation in non-language cortex was also no different from controls.

**Table 2.** Model estimates for whole-brain comparison of people with aphasia and controls 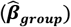 with standard error, and t statistic. The table shows each tissue type at each distance from the lesion (grouped by rows) for the whole-brain analysis of people with aphasia versus control task-relevant activation. The effect size and *P* value is given for the main effect of interest, which is the fixed effect of aphasia status.

| Tissue Type | Distance from Lesion (mm) | fMRI Task |  |  |  |
| --- | --- | --- | --- | --- | --- |
|  |  | Naming |  | Semantic Decision |  |
| | | $\hat{\beta}_{group}$ (SE) | $t(df)$, $p$ value | $\hat{\beta}_{group}$ (SE) | $t(df)$, $p$ value |
| Language | 0-4 | -1.63 (0.28) | $t(1966) = -5.76$, $p < .001$ | -2.51 (0.57) | $t(1105) = -4.37$, $p < .001$ |
| | 4-8 | -1.27 (0.30) | $t(2007) = -4.30$, $p < .001$ | -1.48 (0.62) | $t(1146) = -2.4$, $p = .016$ |
| | 8-12 | -0.98 (0.30) | $t(2048) = -3.29$, $p = .001$ | -0.92 (0.57) | $t(1187) = -1.62$, $p = .105$ |
| | 12-16 | -0.74 (0.32) | $t(1925) = -2.30$, $p = 0.02$ | -0.53 (0.53) | $t(1269) = -0.99$, $p = .324$ |
| | >16 | -0.47 (0.30) | $t(2130) = -1.59$, $p = .111$ | -0.61 (0.46) | $t(1310) = -1.33$, $p = .185$ |
| Language Capable | 0-4 | -1.24 (0.21) | $t(2089) = -5.84$, $p < .001$ | -0.96 (0.41) | $t(1228) = -2.33$, $p = .019$ |
| | 4-8 | -0.90 (0.23) | $t(2089) = -3.98$, $p < .001$ | -0.56 (0.38) | $t(1269) = -1.46$, $p = .144$ |
| | 8-12 | -0.65 (0.23) | $t(2130) = -2.78$, $p = .005$ | -0.54 (0.37) | $t(1269) = -1.45$, $p = .146$ |
| | 12-16 | -0.50 (0.23) | $t(2130) = -2.17$, $p = 0.03$ | -0.46 (0.37) | $t(1269) = -1.27$, $p = .203$ |
| | >16 | -0.23 (0.22) | $t(2130) = -1.07$, $p = .283$ | -0.32 (0.31) | $t(1310) = -1.04$, $p = .297$ |
| Non-language | 0-4 | -0.45 (0.11) | $t(2130) = -4.04$, $p < .001$ | 0.27 (0.19) | $t(1310) = 1.38$, $p = .165$ |
| | 4-8 | -0.27 (0.11) | $t(2130) = -2.48$, $p = .013$ | 0.31 (0.20) | $t(1310) = 1.59$, $p = .110$ |
| | 8-12 | -0.12 (0.11) | $t(2130) = -1.12$, $p = .261$ | 0.21 (0.20) | $t(1310) = 1.03$, $p = .301$ |
| | 12-16 | -0.05 (0.12) | $t(2130) = -0.45$, $p = .656$ | 0.13 (0.20) | $t(1310) = 0.65$, $p = .516$ |
| | >16 | -0.14 (0.10) | $t(2130) = -1.48$, $p = .140$ | 0.06 (0.14) | $t(1310) = 0.47$, $p = .639$ |

**Figure 3.**
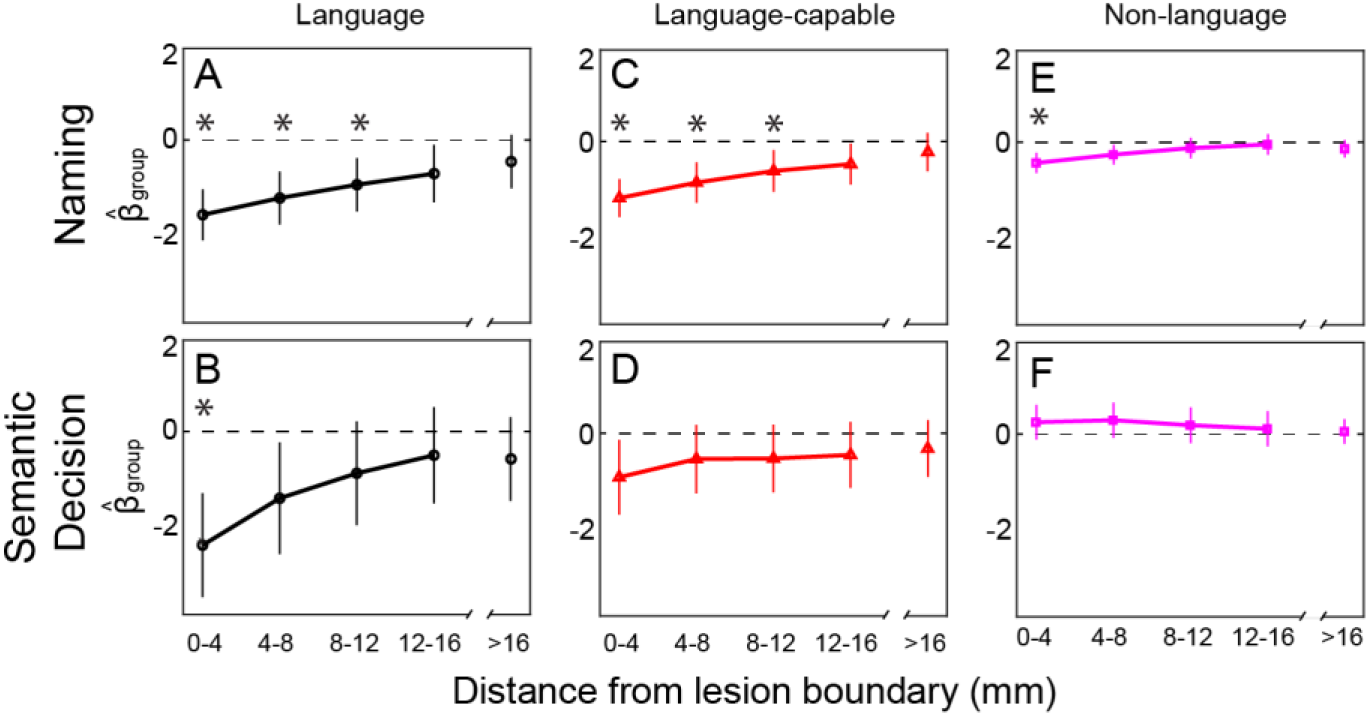
Results from models of effect of group (aphasia vs control) on brainwide activation by tissue type and distance from lesion (*x* axes) for the naming task (top row) and for the semantic decision task (bottom row). The y axis shows estimate of effect of group status 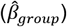 and 95% confidence interval. An asterisk (*) indicates a significant difference between people with aphasia and controls (*P* < .01). The discontinuity in the *x* axes indicates that the rightmost data points included all voxels beyond the perilesional shell. Results are shown for language cortex (A, B) on the left, (C, D) language-capable cortex in the middle, and (E, F) non-language cortex.

### No brain regions exhibit selectively increased activity in perilesional cortex

Although we found no evidence for perilesional plasticity above and beyond typical activation levels in controls when examining hemisphere-wide tissue types, it remains possible that recruitment of perilesional tissue occurs only in certain cortical areas. To assess this, we next compared perilesional activity for people with aphasia vs. controls in individual brain regions defined based on a parcellation atlas (Shen et al., 2013). In the naming task, there were no brain regions in which perilesional tissue exhibited increased activity, but there were several regions with reduced activity in perilesional tissue, including the posterior frontal lobe and operculum, lateral and medial temporal lobe regions, supramarginal gyrus, and occipital cortex (**Fig 4A, Table 3**). In tissue farther from the lesion, increased activity was observed relative to controls in posterior superior frontal sulcus, and reduced activity was observed in lateral occipital lobe and fusiform gyrus **(Fig 4B, Table 3)**.

**Table 3.** Regions of perilesional (4-16mm) and distant (>16mm) abnormal aphasia activation vs control activity during the naming task 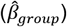 and standard error of the estimate, including the t statistic, degrees of freedom, equivalent P value, and MNI coordinates of the relevant parcel (rows).

| Distance from lesion | Parcel Region | $\hat{\beta}_{group}$ (SE) | t(df), p value | Center of mass (MNI) |
| --- | --- | --- | --- | --- |
| Near lesion (4-16mm) | Inferior precentral gyrus | -1.44 (0.39) | t(1515) = -3.7, P = .005 | -56, -3, 27 |
|  | Frontal operculum | -1.12 (0.34) | t(1638) = -3.29, P = .01 | -56, -3, 6 |
|  | Posterior middle frontal gyrus | -0.92 (0.23) | t(1474) = -4.09, P = .002 | -46, -1, 49 |
|  | Mid/superior operculum | -0.72 (0.25) | t(1351) = -2.87, P = .03 | -39, 9, -5 |
|  | Sup. supramarginal gyrus | -0.74 (0.22) | t(1351) = -3.32, P = .013 | -52, -43, 39 |
|  | Posterior superior temporal sulcus | -0.81 (0.27) | t(1474) = -3.02, P = .02 | -58, -29, 3 |
|  | Inferior occipital gyrus | -1.45 (0.32) | t(695) = -4.51, P < .001 | -41, -51, -17 |
|  | Lateral occipital gyrus | -2.57 (0.68) | t(490) = -3.8, P = .005 | -32, -86, 14 |
|  | Post fusiform gyrus | -2.14 (0.64) | t(449) = -3.33, P = .01 | -44, -70, -14 |
|  | Posterolateral inferior occipital lobe | -1.85 (0.61) | t(531) = -3.02, P = .02 | -35, -84, -7 |
|  | Occipital pole | -2.67 (0.91) | t(285) = -2.93, P = .03 | -15, -97, 7 |
|  | Hippocampus- | -0.82 (0.29) | t(408) = -2.84, P = .03 | -22, -37, 7 |
|  | Hippocampus (mid) | -0.61 (0.19) | t(1187) = -3.19, P = .02 | -36, -24, -15 |
| Distant from lesion (>16mm) | Posterior SFS | 0.69 (0.18) | t(1474) = 3.92, P = .009 | -29, -10, 55 |
|  | Lateral occipital cortex | -1.65 (0.47) | t(1925) = -3.55, P = .01 | -32, -86, 14 |
|  | Posterior fusiform cortex | -1.32 (0.37) | t(2130) = -3.56, P = .01 | -44, -70, -14 |
|  | Posterolateral occipital cortex | -1.41 (0.46) | t(2048) = -3.10, P = .05 | -35, -84, -7 |

**Figure 4.**
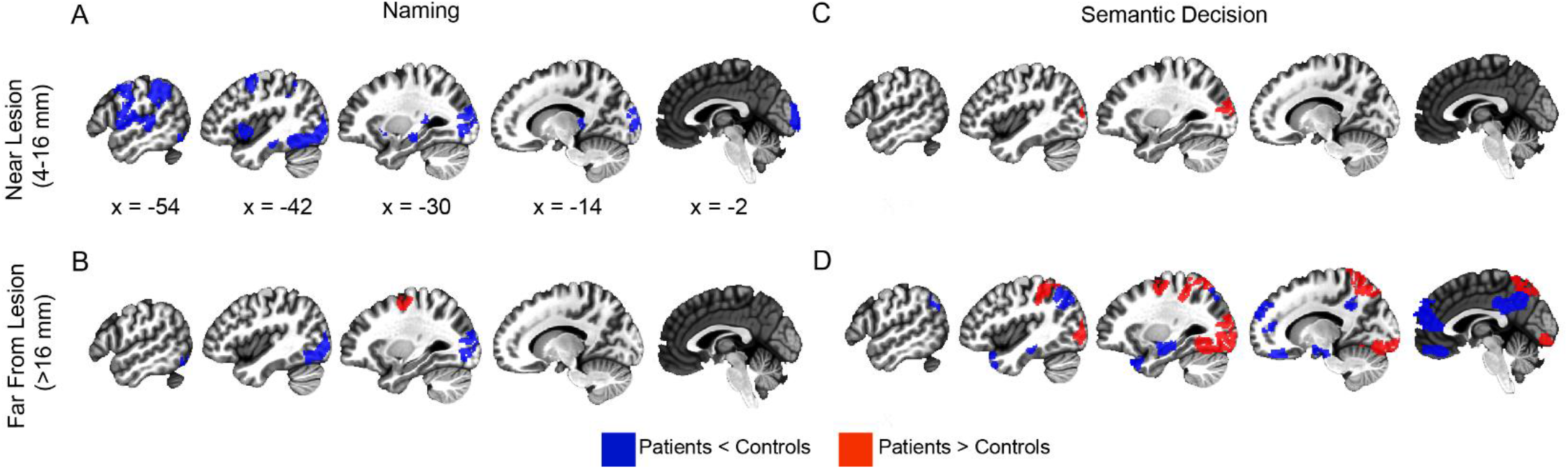
Regional differences in aphasia vs control activity on two language mapping tasks, including naming (left) and semantic decision (right). Regional differences between people with aphasia and controls are shown separately for perilesional tissue (4-16mm of the lesion boundary, top) and for tissue distant from the lesion (>16mm from the lesion boundary, bottom). Blue parcels are controls > people with aphasia, and red parcels are people with aphasia > controls, *P* < .05, Benjamini–Hochberg FDR.

In the semantic decision task, people with aphasia exhibited greater activation than controls in perilesional tissue within lateral occipital cortex, but no systematic decreased activation in perilesional tissue (**Fig 4C, Table 4**). In tissue farther from the lesion, increased activity was also observed in lateral occipital cortex (**Fig 4D, Table 4**), demonstrating that stroke-related increases in activity in this region occurred irrespective of proximity to the lesion. Increased activity was also observed in non-perilesional tissue within the posterior superior frontal sulcus, superior parietal lobule, intraparietal sulcus, and much of the inferior occipital lobe. Decreased activation was observed in tissue distant from the lesion within midline structures such as ventromedial prefrontal and retrosplenial cortices, areas of lateral and medial temporal lobe, and angular gyrus.

**Table 4.** Regions of perilesional (4-16mm) and distant (>16mm) abnormal aphasia activation vs control activity during the semantic decision task 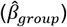 and standard error of the estimate, including the t statistic, degrees of freedom, equivalent P value, and MNI coordinates of the relevant parcel (rows).

| Distance from lesion | Parcel Region | $\hat{\beta}_{group}$ (SE) | $t(df)$, $p$ value | Center of mass (MNI) |
| --- | --- | --- | --- | --- |
| Near lesion (4-16mm) | Lateral occipital cortex | 3.43 (0.97) | $t(326) = 3.53$, $P = .037$ | -32, -86, 14 |
| Distant from lesion (>16mm) | Perirhinal cortex (anterior) | -0.84 (0.30) | $t(1310) = -2.82$, $P = .03$ | -10, 41, -22 |
| | Ventromedial prefrontal cortex | -1.16 (0.37) | $t(1310) = -3.18$, $P = .02$ | -8, 46, 15 |
| | Anterior superior frontal gyrus | -2.04 (0.59) | $t(1228) = -3.47$, $P = .01$ | -12, 56, 29 |
| | Posterior superior frontal sulcus | 1.37 (0.44) | $t(1105) = 3.08$, $P = 0.02$ | -29, -10, 55 |
| | Superior parietal lobule | 1.57 (0.50) | $t(1228) = 3.12$, $P = .02$ | -27, -55, 66 |
| | Medial superior parietal lobe | 1.90 (0.64) | $t(1269) = 3.00$, $P = .02$ | -13, -66, 60 |
| | Intraparietal sulcus | 1.78 (0.51) | $t(1023) = 3.52$, $P = .01$ | -35, -39, 47 |
| | Angular gyrus | -1.60 (0.53) | $t(1064) = -3.05$, $P = .02$ | -41, -67, 42 |
| | Mid anterior temporal pole | -0.90 (0.28) | $t(1269) = -3.17$, $P = .02$ | -33, 18, -33 |
| | Lateral occipital cortex | 1.89 (0.60) | $t(1146) = 3.16$, $P = .02$ | -32, -86, 14 |
| | Medial fusiform gyrus | 1.81 (0.36) | $t(1269) = 5.00$, $P < .001$ | -25, -64, -12 |
| | Posterolateral inferior occipital lobe | 1.84 (0.57) | $t(1187) = 3.24$, $P = .02$ | -35, -84, -7 |
| | Inferior occipital lobe | 1.18 (0.38) | $t(1269) = 3.12$, $P = .02$ | -17, -84, -12 |
| | Occipital pole | 1.92 (0.51) | $t(1269) = 3.78$, $P = .008$ | -20, -96, -9 |
| | Medial suprasplenial cortex | -0.97 (0.35) | $t(1269) = -2.80$, $P = .03$ | -6, -36, 32 |
| | Retrosplenial cortex | -1.61 (0.46) | $t(1310) = -3.52$, $P = .01$ | -8, -55, 37 |
| | Hippocampal formation (anterior) | -0.58 (0.19) | $t(1310) = -2.97$, $P = .02$ | -23, -14, -18 |
| | Hippocampal formation (mid) | -0.57 (0.21) | $t(1310) = -2.65$, $P = .05$ | -36, -24, -15 |

### Perilesional recruitment does not depend on lesion size

Models of aphasia recovery suggest that perilesional recruitment may be particularly evident around smaller lesions. We hypothesized that if smaller lesions were predisposed to perilesional recruitment, then lesion size would correlate with activation in people with aphasia, and individuals with the smallest lesions (<50cc) would exhibit perilesional activation that exceeds the control cohort in the same location.

In the naming task, lesion volume and activity were significantly inversely related for language-cortex both in perilesional tissue (**Fig 5A**) and in tissue far from the lesion (**Fig 5B**). The same pattern was observed in language-capable cortex both in perilesional tissue (**Fig 5C**) and in tissue far from the lesion (**Fig 5D**). There was no significant relationship evident in non-language cortex, whether in perilesional tissue (**Fig 5E**) or in tissue far from the lesion (**Fig 5F**).

**Figure 5.**
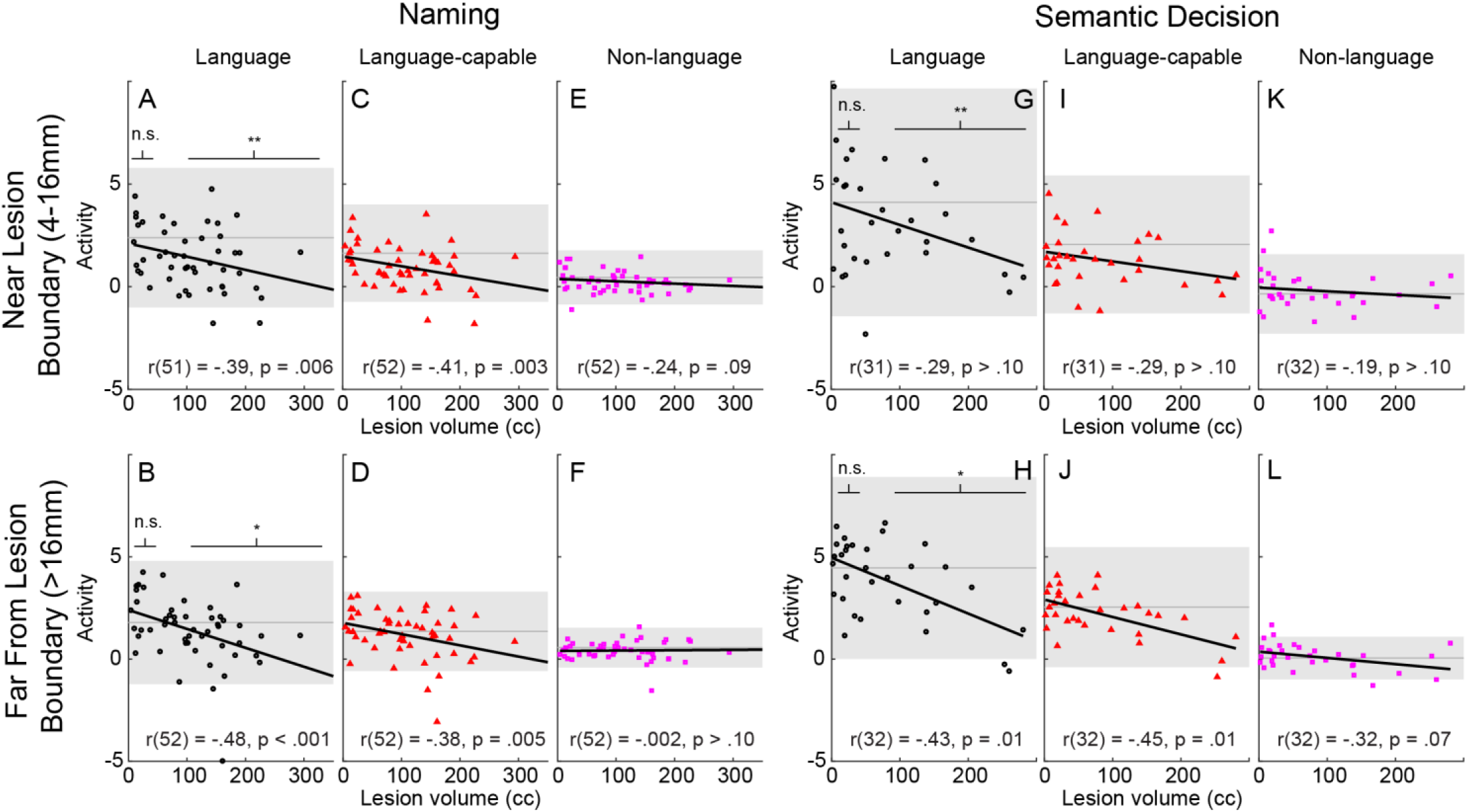
Scatterplots of, lines of best fit for, and correlation coefficients between activation and lesion volume in: perilesional cortex (4-12mm from lesion boundary, top rows) and cortex far from the lesion (>16mm from lesion boundary, bottom rows) for the naming task (left) and semantic decision task (right). Scatterplots are shown for language cortex (black circles), language-capable cortex (red triangles), and non-language cortex (magenta squares). The y axis is the average *t* statistic for the task-contrast within the relevant mask, with each marker representing a single participant with aphasia. The x axis represents lesion volume in cubic centimeters (cc). The mean activation for control subjects is shown as a dark gray line, with the 95% confidence interval shown as a light gray band. Results of LMEM of the effect of group (aphasia vs control) on activity in language tissue are also shown for small lesions (<50cc) and large lesions (>100cc). n.s. = not significant, * = P < .05, ** = P < .001

In the semantic decision task, lesion volume was not related to activity in language-cortex near the lesion, but was inversely related to activity far from the lesion (**Fig 5H**). The same pattern was observed for language-capable cortex (**Fig 5I, J**). There was no significant relationship evident in non-language cortex, whether in perilesional tissue (**Fig 5K**) or far from the lesion (**Fig 5L**).

Although we didn’t observe individuals with increased language activity compared to the control range (**Fig 5, gray band**), there were also not many individuals with decreased activity compared to the control range. Although there weren’t dramatic increases or decreases in activity in individuals, there still might be group effects on average in people with small lesions or people with large lesions, compared to controls. To test this, we broke out a group with small lesions (<50cc) and large lesions (>100cc) to perform a between-group comparison with controls. In both tasks, individuals with small lesions (<50cc) exhibited activity no different from controls in language cortex near or far from the lesion (**Table 5**). In contrast, in both tasks, individuals with larger lesions (>100cc) exhibited significantly decreased activity in language cortex irrespective of distance from the lesion.

**Table 5.**
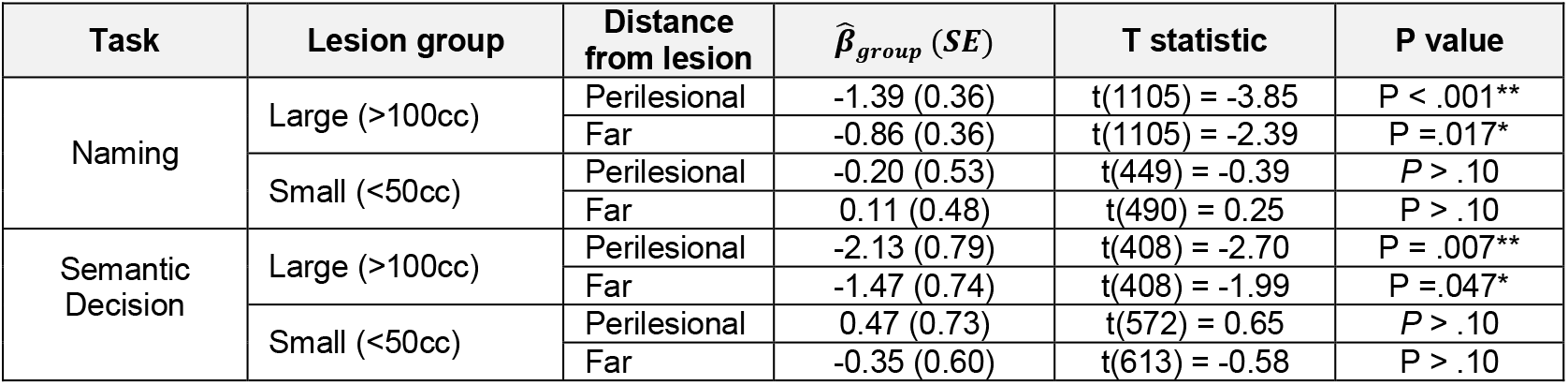
Linear mixed model results of effect of lesion size (large > 100cc, small <50cc) on activity in language tissue, relative to controls, in the naming task and semantic decision task. Results are shown for activity near the lesion (perilesional) and far from the lesion. Results include both the effect of group (aphasia vs control) and its standard error, the t statistic, and p value. ** indicates P < .01, * indicates P < .05

### Disrupted language activity accounts for behavioral impairment

While we didn’t find a relationship between lesion size and perilesional recruitment, we did find that large lesions cause widely disrupted language activity both near and far from the lesion. To address whether these reductions in language activity have behavioral relevance, we next asked whether language activity relates to aphasia severity. In the semantic decision task, there was a significant relationship between activity in task-specific language cortex and overall aphasia severity (WAB AQ), independent of lesion volume, regardless of whether the tissue was perilesional (4-16 mm from the lesion; *r*(31) = .45, *P* = .01) or far from the lesion (>16 mm from the lesion; *r*(32) =.45, *P* = .01). In the naming task, no significant relationship was observed for WAB AQ (perilesional: *r*(50) =.22, *P* > .10; far from lesion: *r*(51) =.17, *P* > .10), or for naming (WAB Naming & Word Finding subscore; perilesional: *r*(51) = .19, *P* > .10; far from lesion: *r*(52) =.20, *P* > .10).

## Discussion

The main goal of this study was to test predictions of the perilesional plasticity hypothesis in post-stroke aphasia. We predicted that recruitment of perilesional tissue through neuroplasticity would result in supranormal task-related activity around lesions. However, we found a brainwide pattern consistent with reduced perilesional activity relative to controls. Moreover, we observed no specific brain regions in which recruitment was evident only near the lesion. When we examined whether perilesional recruitment was evident around small lesions, we found that, although larger lesions were associated with less activity, smaller lesions exhibited perilesional activity no different than controls. Overall, our results are inconsistent with the theory that perilesional plasticity results in recruitment of new brain regions into the language network, or that it results in engagement of typical language regions beyond their normal role in neurotypical individuals.

### Dysfunction and partial normalization of the typical left hemisphere language network

Our results support an alternative interpretation of perilesional recruitment, that strokes to the language network produce network-wide disruptions with decreased language activity, and that perilesional activation is actually just normal activation of unlesioned language processors. We found that the degree of network disruption depended on lesion size, such that large lesions caused widespread disruption, but small lesions resulted in activity no different than controls. Moreover, we found that less disruption of signal in residual language tissue, when measured with a task that produces reliable single-subject maps (Wilson et al., 2018), relates to better behavioral performance even after accounting for the amount of anatomical damage caused by the lesion.

These findings are consistent with previous aphasia treatment studies that found that increased activation in the left hemisphere was associated with improved naming after anomia treatment, with greater increase in activation associated with more improvement (Fridriksson, 2010; Fridriksson et al., 2012; Meinzer et al., 2008) and cross-sectional findings that greater activity in preserved left-hemisphere, relative to controls, was associated with better picture naming performance (Fridriksson et al., 2010). These cross-sectional chronic results also complement studies of spontaneous aphasia recovery that found a recovery trajectory in which good outcomes in the chronic phase were correlated with task-related activity returning to normal levels (Saur, 2006; Stockert et al., 2020). More broadly, these findings are consistent with a recent review of aphasia recovery, which found that lesions caused overall reduced activation in people with aphasia, with activity in left-hemisphere language regions relating to better language function (Wilson & Schneck, 2021).

### Recruitment of alternate left hemisphere processors

In addition to normalization of language processing, previous reports of perilesional plasticity may also reflect increased engagement of alternative left-hemisphere processors irrespective of their proximity to the lesion. This is supported by the regional analysis finding that certain processors were engaged above control levels, but that in every case, these were either regions distant from the lesion or regions that were recruited irrespective of their proximity to the lesion.

Several types of processes might underlie the recruitment we measured as increases in alternative left-hemisphere processors. For instance, the increased activation might relate to compensatory plasticity (Takeuchi & Izumi, 2013), the use of compensatory strategies relying on spared ability (Bury & Jones, 2002), or network-specific changes such as increased reliance on “domain general” processes (Geranmayeh et al., 2014). Our finding of increased activity in posterior superior frontal lobe and parietal lobe shows consistent localization with a domain general dorsal attention/salience network (Fedorenko et al., 2013). Previous work has found increased left-hemisphere activity in people with aphasia during language processing, but a common region exhibiting increased activation would be unlikely to be perilesional since perilesional tissue would be in different places for different individuals (Brownsett et al., 2014; Geranmayeh et al., 2014). Thus, greater activation observed in these regions might relate to compensatory increased reliance on domain-general processing for language tasks. Our finding of increased activity in lateral occipital cortex, irrespective of distance from the lesion, might be explained by recruitment of additional visual processing of written stimuli in the semantic decision task.

### Task-independent and task-dependent effects

One limitation of prior fMRI studies of aphasia is that they have typically examined only one task in a single group of participants. Much of the heterogeneity of results in the literature likely results from idiosyncrasies of individual samples of participants or the tasks used to elicit language activity. Here, we compared results from the same analysis approach using two different tasks in two different groups of participants. The results addressing the question of perilesional plasticity are remarkably consistent across the two tasks, providing very strong support for the conclusions above. However, there were some different findings between tasks. Not surprisingly, different regions were engaged by the two tasks, and therefore the localization of effects in the regionwise analysis was different. Additionally, the behavioral relationships are stronger for the more reliable task (although they numerically trend in the same direction in the naming task, they don’t approach significance). The stronger relationship with behavior supports the use of reliable tasks for questions related to neuroplasticity (Wilson et al., 2019).

### Limitations and alternative interpretations

One limitation of this study is that we did not characterize potential perilesional hypoperfusion. However, we observed effects outside the 8mm range of hypoperfusion measured by Richardson et al. (2011). We also observed reductions in activity distant from the lesions and increased activity in perilesional tissue in one region in the semantic decision task, with similar levels of increased activity when the tissue was perilesional and when it was not. This strongly suggests that perilesional hypoperfusion was not a major factor in the observed effects. In addition, our participants were all in the chronic stage of recovery, and we did not examine the transition from acute to chronic, or directly assess effects of treatment. Perhaps perilesional plasticity is transiently observable during recovery, or only in a behaviorally-enriched treatment context. Future treatment studies should conduct analyses similar to those presented here to test if increases in perilesional activity extend beyond the typical level of activation in controls.

### Conclusions

In conclusion, we found no evidence for perilesional plasticity measured by BOLD fMRI in two groups of people with chronic aphasia using different tasks. We did find evidence for lesion-size dependent language network dysfunction, suggesting that normalization of task-related activity may explain some of the conclusions drawn from previous studies. These results place constraints on mechanistic accounts of chronic post-stroke aphasia neuroplasticity measured with BOLD fMRI.

## Acknowledgments

The authors would like to thank Mackenzie Fama, Zainab Anbari, and Kate Spiegel for data collection. We would also like to thank the research participants, without whose donated time and efforts we could not have conducted this study.

## Author Contributions

ATD formulated the general research question; ATD and PET conceptualized the analyses, ATD performed all analyses; ATD contributed a draft of the text and figures. CV, SP, ED, EL, and SS contributed recruitment, testing, scoring and interpreting participant scanning and behavioral data. All authors revised and edited the final manuscript.

